# Conformational dynamics of Tau in the cell quantified by an intramolecular FRET biosensor in physiological and pathological context

**DOI:** 10.1101/041756

**Authors:** C Di Primio, V Quercioli, G Siano, B Kovacech, M Novak, A Cattaneo

## Abstract

Impaired interactions of Tau protein with microtubules (MT) and Tau misfolding play a key role in Alzheimer disease (AD) and other neurodegenerative diseases collectively named Tauopathies. However, little is known about the molecular conformational changes that underlie Tau misfolding and aggregation in pathological conditions, due to the difficulty of studying structural aspects of this intrinsically unfolded protein, particularly in the context of living cells.

Here we developed a new Conformational-Sensitive Tau sensor (CST), based on human Tau full length protein, to investigate the changes in 3D conformation and aggregation state of Tau upon modulation of its interactions with MTs in living cells, in physiological and pathological conditions. After showing that the CST fully preserves functional Tau activities in living cells, we demonstrated that MT-bound Tau displays a loop-like conformation, while soluble Tau assumes a relaxed conformation.

The imaging readout based on CST allowed to discover new conformational properties of full length Tau in living cells, when challenged with Alzheimer-relevant seeds from different sources, and to learn about different ways to induce the self-aggregation of full length Tau in cells. Furthermore, it allowed to investigate the contribution to the pathology of point mutations known to alter Tau/MTs interaction.

**SIGNIFICANCE:** The microtubule-associated protein Tau regulates the stability of microtubules (MT) and its alteration and misfolding plays an important role in Alzheimer disease (AD) and in several other neurodegenerative diseases collectively named Tauopathies. However, little is known about the molecular mechanism that modulate the conformational changes of Tau in pathological conditions. Here, we developed a conformational sensitive Tau biosensor providing insights into the 3D conformation of Tau in living cells. This new tool showed that MT-bound Tau displays a loop-like conformation, while soluble Tau assume a relaxed conformation, which might be more prone to aggregation. Furthermore the CST was used to develop an assay to screen molecules and mutations involved in Tau self-aggregation in living cells. Remarkably, the imaging tool developed allowed to demonstrate that Aβ oligomers induced Tau aggregation in living cells.

## INTRODUCTION

The microtubule associated protein Tau plays a central role in maintaining the stability of neuronal cytoskeleton and regulating axonal trafficking. In the adult human brain there are six isoforms of Tau, with three or four C-terminal microtubule-binding repeats, through which they bind microtubules (MTs) and promote tubulin polymerization, thus stabilizing MT network.

Tau misfolding and Impaired interactions with MTs play a key role in Alzheimer disease (AD) and other neurodegenerative diseases collectively named Tauopathies ^1^. In these pathological conditions, Tau self-assembles into insoluble intracellular inclusions, known as neurofibrillary tangles (NFTs). Thus, Tau NFT pathology is a hallmark that links all these disorders to a shared mechanism of neurodegeneration.

Little is known about the molecular conformational changes that underlie Tau misfolding and aggregation, due to the difficulty of studying structural aspects of this intrinsically unfolded protein, particularly in the context of living cells. Understanding the molecular basis of Tau perturbed behavior is essential for the development of effective therapies.

Great efforts have been made to study Tau conformation changes, misfolding and aggregation *in vitro*, using purified proteins^2-6^. However, in the cellular background, Tau behavior and MTs interactions are modulated by Tau post-translational modifications such as phosphorylation, acetylation and truncations ^7-19^ that at different extent induce its detachment from MTs and aggregation into insoluble, filamentous inclusions^20-24^.

Assembled Tau has been shown to behave like a prion, in that assembled Tau can seed the aggregation of native Tau ^25-27^.The seeding activity of recombinant Tau repeat domain (RD) and Tau assemblies, purified from AD or Tauopathy brains, and the cell-to-cell propagation of misfolded aggregated Tau, has been recently studied in cells expressing a biosensor system based on the aggregation-competent core Tau repeat domain ^2,28-33.^However, studying tagged Tau repeat domain (RD) fragments does not allow to investigate the misfolding and aggregation process of full length native Tau nor to take into account cellular cofactors modulating Tau conformation and misfolding outside the RD domain. In other cell models, it was shown that the seeding activity of full length native or recombinant Tau is dependent on its aggregation and that Tau conformation determines the seeding potencies of Tau aggregates^20,34,35^. This underscores the importance of developing tools to monitor Tau conformation in the context of living cells.

Here we describe a new Conformational Sensitive Tau sensor (CST), based on Tau full length protein and we use it to demonstrate that, under physiological conditions, Tau assumes very different 3D conformations in the cell, as a function of its interactions with MTs. The CST reporter can be used to monitor the effects of treatments that reduce the levels of cellular Tau or compete with its interaction with MTs. CST reporter cells have then allowed demonstrating the conformational changes and assembly that Tau undergoes when cells are exposed to proteopathic seeds of different nature, including synthetic Tau seeds, A b oligomers and mouse and human brain extracts. Finally, the CST has been used to study Tau /MTs interaction and aggregation propensity in the cell, in the context of different Tau point mutations linked to FTD or to its phosphorylation.

## RESULTS

### Conformational-Sensitive Tau Sensor preserves functional activities of Tau

To investigate Tau structure and function in living cells we generated a conformational sensitive fluorescent sensor by fusing ECFP at the N-terminus and EYFP at the C-terminus of the full length Tau-D sequence (0N4R)(fig. 1A). A flexible linker (RSIVT) connects Tau to the fluorophores, at either end. It is expected that, depending on the conformation of Tau, the ECFP and the EYFP moieties of the sensor could generate a FRET signal, thus realizing a context-dependent conformational-sensitive Tau sensor (CST). Mono-labeled chimeric constructs have been used as controls. HeLa cells transiently transfected with the CST sensor have been analyzed by western blot (fig. 1B). The intact CST protein showed the expected molecular weight of 120KD. Control blots with the C3 Tau antibody demonstrated a partial cleavage of Tau at Asp421 (Suppl.Fig.1). The same processing also occurs in mono-labeled and unlabeled Tau constructs, confirming that, in HeLa cells, Tau undergoes a partial proteolytic processing^22^. Nevertheless, control blots, with antibodies against the microtubules binding domains (MTBD) and against GFP, showed that CST processing is restricted to a very low amount of protein.

**Figure 1.**
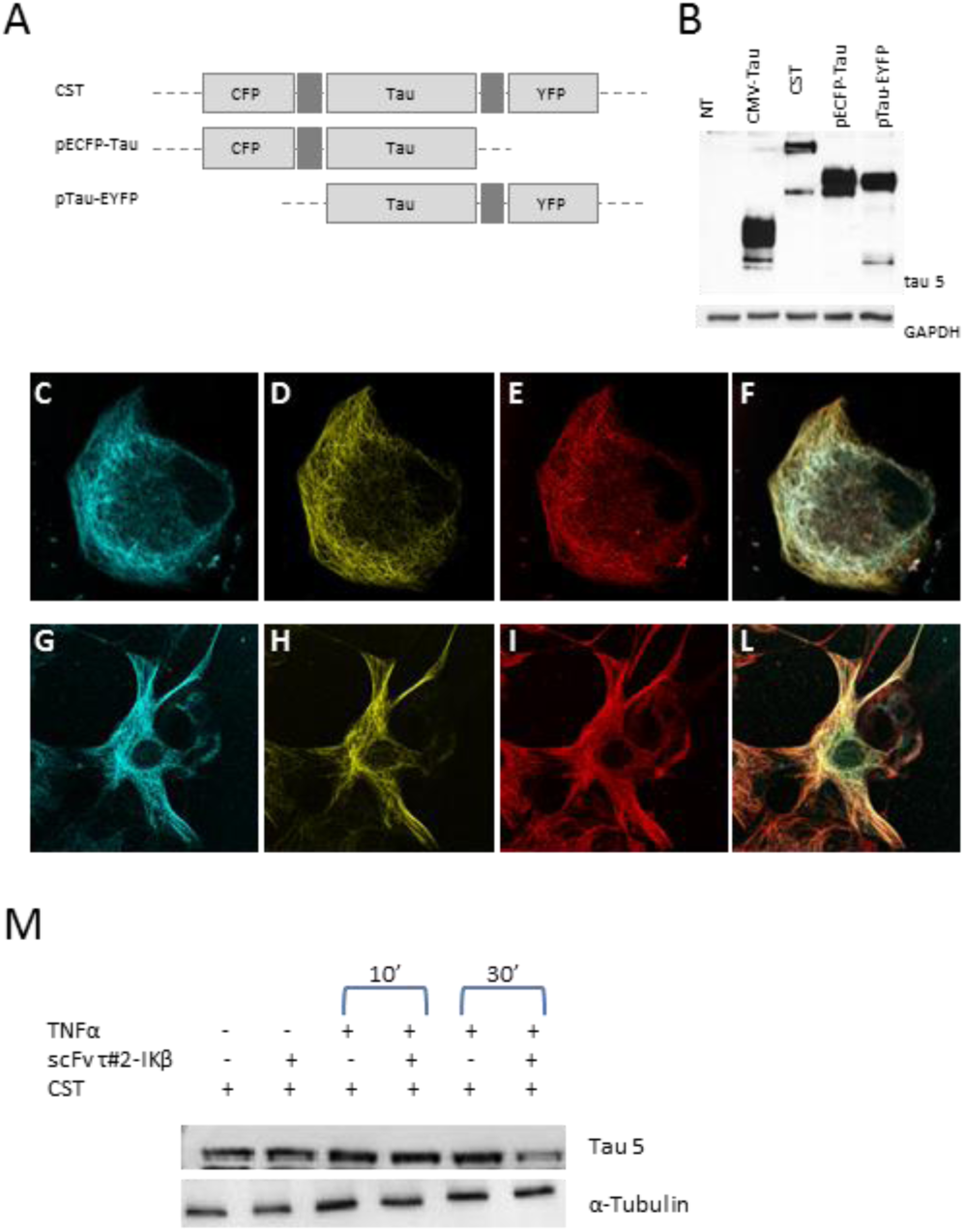
CST molecular and imaging characterization. A. Schematic representation of chimeric constructs. B. Western blot analysis of unlabeled CMV-Tau, CST, ECFP-Tau and Tau-EYFP in total extracts from HeLa cells. C-F. Imaging of HeLa cells co-transfected with CST and tubulin-RFP, ECFP donor channel (blue), EYFP acceptor channel (yellow), tubulin-RFP (red), merge. G-L. Imaging of HT22 cells co-transfected with CST and tubulin-RFP, ECFP donor channel (blue), EYFP acceptor channel (yellow), tubulin-RFP (red), merge. M. CST modulation by the suicide antibody scFv τ#2-IKβ. Western blot of cells transfected with scFv τ#2-IKβ or with CST have been treated or not with TNFα for 10 minutes or for 30 minutes.

Confocal microscopy imaging, in both CFP and YFP channels, showed that CST sensor, as well as mono-labeled and unlabeled Tau, decorated the cellular microtubules network (MTs) and colocalized with the tubulin-RFP, both in HeLa cells and in immortalized hippocampal neurons HT22 (fig. 1C-L; Suppl.Fig. 1C,D,E). Moreover, the CST sensor displayed the same localization of the endogenous Tau, visualized in SH-SY5y neuroblastoma cells by indirect anti tubulin immunofluorescence (Suppl. Fig. 1 D,E).

Notably, overexpression of CST did not result in any detectable toxicity and transfected cells were morphologically intact for throughout the observation time (at least 5 days, data not shown).

Taken together, these results demonstrated that in physiological conditions the CST sensor preserves the basic functions of Tau and its ability to interact with MTs.

### Intracellular CST can be degraded by a suicide anti Tau intrabody

Having established that CST recapitulates the behavior of Tau, we probed its ability to interact in living cells with an anti Tau intrabody^36^. Reducing the levels of cellular Tau is considered a promising strategy to limit the consequences of Tau misfolding and aggregation. To modulate the levels of Tau, we exploited SIT (Suicide Intrabody Technology)^37^, by expressing the anti Tau scFv τ#2 single chain Fv fragment in the format of an inducible suicide intrabody (scFv τ#2-IKP). In this format, treatment with TNFα of the cells expressing the scFv τ#2-IKβ intrabody determines its transition from a long-lived protein into a short lived, rapidly degraded protein, that escorts the bound Tau protein to proteasomes^37^. HeLa cells were co-transfected with the CST sensor and the suicide anti Tau intrabody scFv τ#2-IKβ. The treatment of cells with TNFα for 30 minutes induced the degradation of scFv τ#2-IKβ and eventually the concomitant degradation of targeted Tau protein. We observed that, similarly to endogenous Tau, CST is likewise subjected to this degradation by the scFv τ#2-IKβ (Fig.1 M). These results provide a further validation to the concept of using SIT to reduce the cellular levels of Tau and demonstrate that, in the cellular environment, CST stability can be modulated by scFv τ#2-IKβ, similarly to that of endogenous Tau^37^. Moreover, CST can be used to monitor the outcome of experimental strategies aimed at regulating the overall levels of Tau in a target cell.

### Tau bound to microtubules display a loop-like 3D conformation

Despite the great relevance of Tau conformational changes as determinants for its pathological oligomerization and aggregation^2,5,20,28,38-40^, little is known about these modifications, due to the difficulty of studying structural aspects of this intrinsically unfolded protein, particularly in the context of living cells.

To exploit the CST sensor as a conformational probe for Tau, we used quantitative FRET microscopy for the detection of protein conformational changes in the cell, based on the energy transfer between the ECFP and EYFP fluorophores, when the N-and C-termini of the CST are close enough (<50Å) to allow the process.

The quantification of FRET by sensitized emission has been performed in HeLa cells expressing the CST. Figure 2A shows donor and acceptor imaging of CST and also a normalized FRET image (NFRET)^41^, indicating that CST displayed a FRET-positive signal mostly along MTs. Indeed, along selected lines (white line Fig.2B,C) the MT-bound CST displayed NFRET values between 15-30.

**Figure 2.**
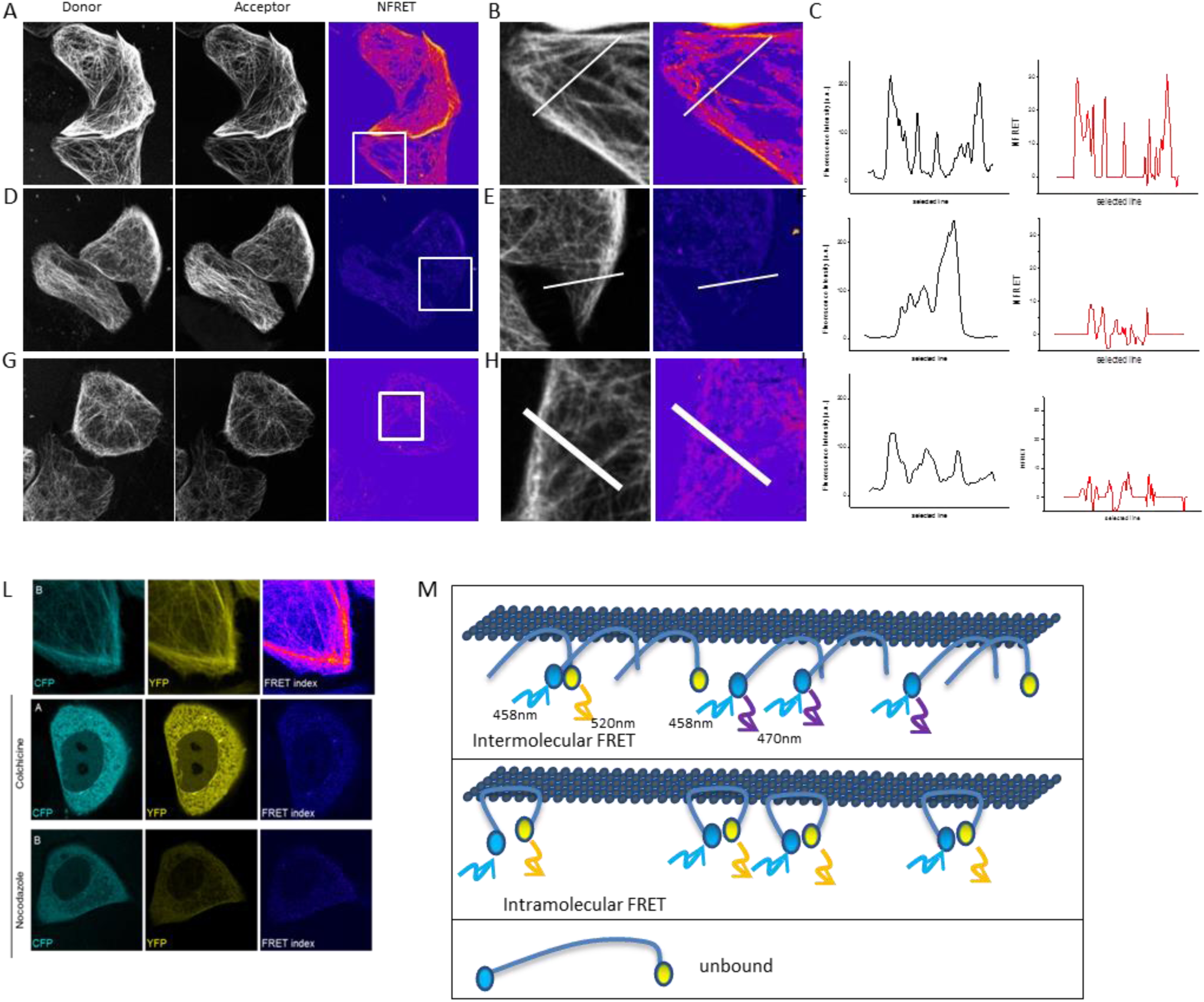
Intramolecular FRET measurements. A. Donor channel (grey), acceptor channel (grey) and NFRET images (magenta) in cells expressing CST. B. Fluorescence intensity image in the outlined region (grey, left panel) and related NFRET image (pink, right panel). C. Fluorescence intensity profile along the white line (left, black graph) and NFRET intensity profile along the white line (right, red graph). D. Donor channel (grey), acceptor channel (grey) and N FRET image (magenta) in cells co-expressing CFP-Tau and Tau-YFP. Imaged cells displayed a fluorescence intensity comparable to CST. E. Fluorescence intensity image in the outlined region (grey, left panel) and related NFRET image (pink, right panel). F. Fluorescence intensity profile along the white line (left, black graph) and NFRET intensity profile along the white line (right, red graph). G. Donor channel (grey), acceptor channel (grey) and N FRET image (magenta) in cells co-expressing CFP-Tau and Tau-YFP. Imaged cells displayed a half fluorescence intensity in comparison to CST. H. Fluorescence intensity image in the outlined region (grey, left panel) and related NFRET image (pink, right panel). I. Fluorescence intensity profile along the white line (left, black graph) and NFRET intensity profile along the white line (right, red graph). L Donor channel (blue), acceptor channel (yellow) and NFRET image (pink) for cells expressing CST, untreated (upper panel) and treated with colchicine (middle panels) and nocodazole (lower panels). M. Intramolecular and intermolecular FRET schematic representation.

To verify whether the FRET signal observed was due to intramolecular or to intermolecular interactions between the Tau N-and C-termini, cells co-expressing monolabeled constructs (ECFP-Tau and Tau-EYFP) have been analyzed. As shown in figure 2, in cells expressing monolabeled constructs with comparable concentrations of fluorophores with respect to CST (fig.2 D-F) or with half concentration of fluorophores (G-I) the NFRET value is around 10. These controls allowed to verify the FRET values in the presence of double Tau concentration or the same moiety with respect to CST, respectively (Suppl fig. 2). Given the small distances involved in FRET process, these results indicate that MT-bound CST assumes a loop-like 3D conformation, with its N-terminal and C-terminal domains separated by a distance shorter than 50Å, allowing FRET to occur. On the contrary, in conditions where only intermolecular FRET can occur, such as when monolabeled Tau is expressed, the FRET signal is significantly lower.

To analyze the 3D conformation of soluble CST, not bound to MT, cells were treated with the MT disrupting drugs nocodazole or colchicine. When MTs are disrupted by these drugs, the CST diffused into the cytoplasm (Suppl. fig.3). Quantitative analysis revealed that cytoplasmic CST did not display FRET (fig.2 L). Western blot analysis showed that, in the absence of MTs, most of the CST molecules are intact (Suppl.fig.3), thus the lack of FRET signal is not due to the cleavage of fluorophore moieties. These results indicate that Tau molecules not bound to MTs assume a more relaxed conformation that does not allow the energy transfer between its C-and N-termini, confirming our conclusion that the FRET signal is intramolecular and peculiar of intact CST bound to MTs (fig.2 M).

### CST-based assay detects molecules potentially involved in pathological Tau self-aggregation

Current views indicate that misfolded Tau drives its aggregation and spreading in Tauopathy disease progression^6,22,25,34,42,43.^ We decided therefore to exploit the CST sensor to detect the seeding induced by MTs interfering molecules and agents potentially involved in Tau self-aggregation.

To this aim, CST reporter cells were first exposed to extracellular synthetic Tau seeds, prepared from (297-391)4R recombinant Tau fragment^3,4^ as a positive control for Tau seeding activity. As expected, 48 hours later the Tau network was completely displaced from MTs and Tau cytoplasmic inclusions appeared (Fig. 3). Remarkably, fluorescent CST aggregates displayed both CFP and YFP signals. We then challenged the CST reporter cells with molecules involved in MTs dynamic modulation, the anti-mitotic agents paclitaxel (PTX) and its derivative epothilone D (EpoD). Both drugs promote the assembly of tubulin and MTs stabilization and share a common binding site on tubulin with Tau. We observed that soon after the PTX addition the CST detached from the stabilized MTs and relocalize into fluorescent inclusions that are composed by both fluorophores. The inclusions were also positively stained by the amyloid-specific dye K114 (Suppl. Fig.4). Conversely, epothilone D caused only a mild relocalization.

**Figure 3.**
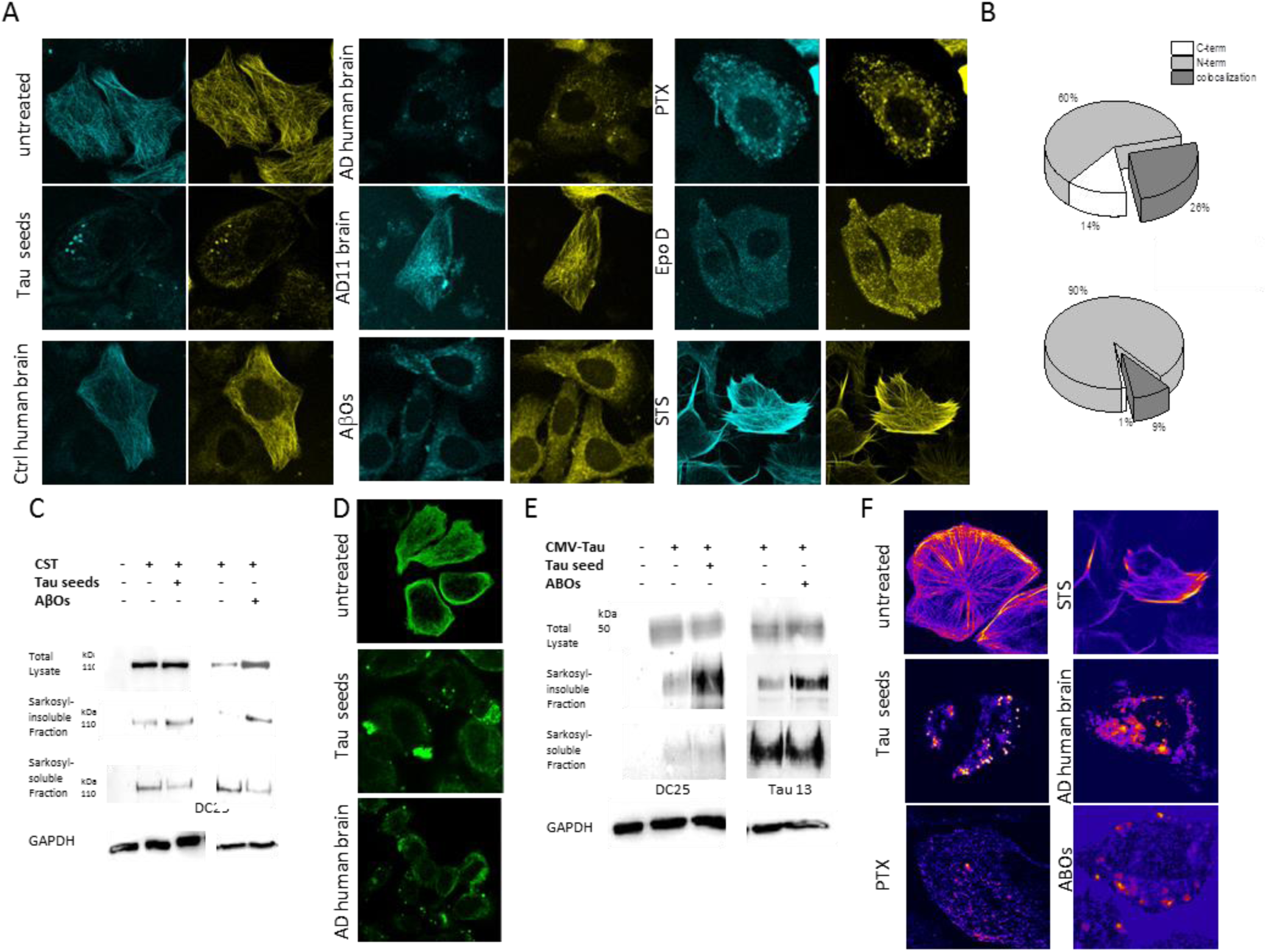
CST imaging in cells treated with different agents involved in self-aggregation. A. Control and treated cells have been imaged in both CFP and YFP channel in living cells. CST reporter cells have been treated for 48h with Tau seeds 200nM, for 96h with 10ug of control human brain lysate or AD human brain lysate, with AβOs or with PTX 1uM or EpoD 1uM. B. Quantification of fluorescent aggregates detected in reporter cells treated with AD human brain lysates (upper graph) or with ABOs (lower graph). C. Western blot of Sarkosyl soluble and insoluble fractions of reporter cells treated with Tau seeds or AβOs. D. Tau imaging in cells treated with Tau seeds (middle panel) and AβOs (lower panel). Cells expressing unlabeled Tau stained with anti Tau13 antibody (green). E. Western blot of Sarkosyl soluble and insoluble fractions of cells expressing unlabeled Tau and treated with Tau seeds or AβOs. F. FRET images of untreated and treated reporter cells.

Tau phosphorylation state modulates its interaction with MTs, due to the repulsive action of positive charges. To investigate the ability of CST in detecting this modulation, CST reporter cells were treated with Staurosporin (STS), a drug that inhibit kinases and could modify Tau propensity to bind MTs. This treatment resulted in a stronger CST signal on MTs indicating that the impaired phosphorylation of Tau increased the interaction between Tau and MTs (fig.3 A).

CST-reporter cells were then exposed to extracts from AD and control human brains (n=2 each). Figure 3 shows that cleared lysates from AD brain, but not from control brains, induced a significant relocalization of Tau away from MTs, forming intracellular inclusions. Remarkably, the MT network is not grossly altered by Tau displacement (suppl. Fig.5) and we observed that while 26% of the inclusions displayed both fluorophore signals, the majority displayed only the N-terminal signal (ECFP labeled) and 14% contained only the C-terminus (EYFP labeled) (Fig. 3 B). Thus, the CST detects an unequal accumulation of processed aggregation Tau protein seeded the AD brain lysate.

CST reporter cells were also challenged with brain cleared lysates from the anti NGF AD11 mouse model^44,45^ (that develop a progressive Alzheimer-like neurodegeneration) and from age matched control mice. As with AD human brain lysates, we found that adding AD11 lysates induced Tau relocalization and aggregation, with the prevalence of N-terminal ECFP positive inclusions.

AD brain lysates contain many different components that might potentially act as seeds for Tau aggregation, including other misfolded proteins such as the Aβ peptide. In order to verify whether the observed seeding is specific for Tau, or whether reporter cells are also sensitive to other factors, we probed CST reporter cells with preparations containing natural Aβ oligomers (AβOs). We used AβOs produced by the 7PA2 fAD cell line (CHO cells overexpressing fAD human V717F APP mutant), whose supernatants are considered the best characterized source of natural, pathologically relevant AβOs^46^. Surprisingly, AβO-containing 7PA2 supernatant induced a significant relocalization of Tau, with the appearance of numerous inclusions with the prevalence of CFP positive spots (fig.3A, B). These data have been confirmed also in HT22 cells (Suppl. Fig.6) and are not observed when either cell is exposed to control CHO supernatant.

In order to verify that the observed CST-positive inclusions do contain aggregated Tau, we performed Sarkosyl fractionation of the CST reporter cells treated with Tau seeds and ABOs. As shown in Fig. 3C, western blot analysis of the Sarkosyl extracts with the anti Tau mAb DC25^47^detected the significant presence of full length CST into the Sarkosyl-insoluble fraction of both treated cells, with a concomitant decrease of CST in the soluble fraction.

Taken together, these results indicate that full length Tau undergoes to aggregation in response not only to homotypic seeding, by Tau seeds or by brain extracts containing Tau seeds, but also in response to heterotypic seeding, by cell supernatants containing AβOs, but no Tau seed. To further verify that the pathological interference of AβOs on full length Tau aggregation was not due to the peculiar architecture of CST, we challenged cells expressing unlabeled Tau. The immunofluorescence for Tau in control and ABOs-or Tau-treated cells (figure 3.D) confirmed that AβOs induced Tau relocalization and aggregation as effectively as Tau seeds. Further analysis of sarkosyl-insoluble and soluble fractions showed that full length Tau accumulates into insoluble fractions in both conditions, confirming the ability of AβOs to induce Tau aggregation (figure 3.E) and further confirming that the fluorophores into the CST construct do not alter the native conformational changes that Tau undergoes during neurodegeneration.

Furthermore we investigated whether the CST could detect by FRET conformational changes occurring during Tau aggregation. We observed that in cells, treated with Tau seeds or with PTX, FRET signal is undetectable on MTs, while aggregates displayed a positive signal. Moreover, FRET signal persists on MTs after STS treatment (Fig 3 F). FRET signal from cytoplasmic inclusion has been confirmed by the same FRET experiment performed in cells expressing CFP-TAU and TAU-YFP constructs and probed with Tau seeds (data not shown).

Taken together, these results indicate that Tau undergoes aggregation in cells, in response to synthetic Tau seeds as well as to complex lysates derived from brains of neurodegeneration mouse models or Alzheimer's disease patients, producing an unequal assembly of N-terminal cleaved species. Heterotypic seeding by AβOs was also demonstrated. Remarkably, Tau aggregation occurs also after its massive displacement form MTs by paclitaxel, while phosphorylation inhibition strengthens Tau/MTs interaction.

### CST-based assays in the context of different Tau point mutations

The mechanisms by which Tau mutations induce pathology are diverse and not always clear. Evidence supports changes in alternative splicing, phosphorylation state, interaction with tubulin, and self-association into filaments as important contributing factors ^48-50^. The lack of suitable tools to investigate the contribution of individual mutations to Tau pathology at the cellular level prompted us to set up an assay based on mutated CST sensor to investigate the mobility and the conformation of Tau monomers.

Tau mutations known to alter the interaction with MTs or to promote self-aggregation^24,28,34,51-55^ have been inserted into the CST constructs. P301S or P310L mutations determine accelerated filament formation *in vitro* and NFTs development in transgenic mice^56-59^. Furthermore, the phosphorylation of S422 is considered an early biomarker of AD progression and the pseudo-phosphorylated form S422E, but not the S422A unphosphorylated form, has been reported to have a pro-aggregation property^60^.The phosphorylation of AT8 epitope (S199/S202/T205) has been reported as a late biomarker of AD ^61^. Reporter cells expressing CST bearing these single mutations have been subjected to imaging, FRET and FRAP analysis. We first observed that the AT8 epitope is constitutively phosphorylated in Hela cells (Data not shown).

We then investigated the mobility of Tau molecules in cells, by FRAP experiments. A selected region of interest in cells expressing Tau mutation bearing CST was photo bleached and the recovery rate of the fluorescence intensity, due to the mobility of neighboring fluorescent molecules, was evaluated. FRAP analysis showed that 73% of the CST molecules are in the mobile fraction, contributing to fill the bleached area. The recovery curve analysis revealed that two main components are represented: the 57% ascribed to a MTs-bound phase that is characterized by a τ_1_ of 30 sec and the 16%, ascribed to diffusing soluble molecules with a τ_2_ of 4.3 sec.

The same FRAP analysis was performed in reporter cells expressing Tau mutation-bearing CST sensors. We observed that S422A-CST and AT8mim-CST displayed the same recovery curves of wild type CST (fig. 3.B). On the contrary, P301L-CST, P301S-CST, S422E-CST displayed a higher mobility, since the mobile fraction increased respectively to 84%, 85% and 86% due to an increase of both the diffusive and the bound fractions. In particular, the P301L mutation induced a remarkable increase in the diffusive fraction (38%). These results suggested that these mutations weaken Tau interaction with MTs (fig. 3B, Suppl. Table 1). Conversely, S199A/S202A/T205A mutations abolishing AT8 phosphorylation epitope resulted in a slower mobility of Tau. Indeed, the mobile fraction decreased to 57% and the τ_2_ increased to 46.8 indicating that the protein is strongly attached to MTs.

Altogether the FRAP results indicate that tested mutations cluster into three groups, according to their impact on the CST behavior in cells: AT8mim and S422A do not alter the mobility of Tau, P301L, P301S and 422E altered Tau/MTs interaction, conferring an increased mobility, while the AT8 null mutation strengthens Tau interaction with MTs.

FRET results on MTs of CST reporter cells fit with these three clusters (fig.3 C). Indeed, the increased FRET signal observed with the AT8 null mutation could indicate a conformational change of Tau that strongly binds MTs. On the other hand, the reduced MT FRET signal observed for the P301L-CST could be due to protein conformational changes interfering with MTs interactions.

CST reporter cells for the P301L and the AT8 null mutations were subjected also to imaging analysis. We observed that while the decoration of MT network by CST is totally superimposable to that by anti-tubulin antibodies, P301L-CST undergoes to a partial relocalization in the cytoplasm and a breakdown in the extent of network decoration, that was morphologically quantified by measuring the total filament length (summing piecewise filament segments) and the number of branching points (see methods) (fig. 3D). Image analysis demonstrated for both parameters a 2 fold decrease of P301L-CST, with respect to CST. On the contrary, AT8mut-CST displayed an increased Tau network complexity.

## DISCUSSION

The MT binding Tau protein is an intrinsically unfolded protein^6,15,35,39,62^. Despite the importance of the regulation of Tau/microtubules interaction, little is known about the conformational changes to which Tau is subjected between the MT-bound and unbound states, both in physiology and pathology ^63^. No direct information is available on the molecular mechanism involved on these conformational changes in the cell.

The CST developed in our study allowed to show that the MT-associated Tau displays a loop-like 3D conformation, in the live cell, with its N-and C-termini close to each other, at a distance sufficiently small to allow for a FRET signal to be elicited. This loop structure is lost in MT-unbound molecules, soluble in the cytoplasm.

Having established that the CST recapitulates the main cellular features and properties of Tau, the CST formed the basis of cellular assays to monitor i) the effectiveness of strategies to reduce the cellular levels of Tau, ii) the seeding activity of small molecules, iii) the seeding activity on Tau aggregation by complex mixtures such as AD brain extracts or by heterotypic seeds such as amyloid Aβ oligomers, iv) the impact of different Tauopathy-related Tau mutations or of Tau phosphorylation sites on the interaction of Tau with MTs.

CST-based assays show that synthetic Tau has a significant seeding activity, inducing detachment of Tau from MTs, the formation of Tau inclusions and the partitioning of Tau into the Sarkosyl insoluble fraction, indicating the formation of filamentous aggregated Tau. Interestingly, the Sarkosyl insoluble fraction contained full length Tau molecules, showing that the detachment of Tau from MTs and the formation of inclusions precede its cleavage.

Seeding components present in the AD brain lysates induced a similar result, as might have been expected since they contained also Tau aggregates. However, their heterogeneous and complex composition, containing also fibrillary and oligomeric Aβ open the key question of which is the component in the AD lysates that is responsible for inducing Tau missorting and aggregation. For this reason, we investigated the contribution of Aβ oligomers to the Tau seeding activity. We observed a partial Tau relocalization away from MTs and its aggregation. The mechanism of this heterotypic seeding of Tau aggregation has to be further investigated, since AβOs could act in a direct way after internalization (cross-seeding) or indirectly, via the activation by AβOs of signaling pathways that could impact on Tau mislocalization. Nevertheless, aggregates formed by treating cells with AD brain lysates or ABOs shared a peculiar feature that is the unequal accumulation of Tau fragments. Indeed, while aggregates from synthetic Tau seeds treatment displayed both fluorophores and a positive FRET signal, aggregates formed in response to AβOs or AD brain treatment showed fewer FRET-positive aggregates. This suggest that the seeding activity exerted by synthetic Tau could imply the detachment of intact Tau from MTs and its aggregation, while ABOs and other seeds contained into the brain lysates could activate a proteolytic process that ends in the aggregation of truncated Tau.

Remarkably, the CST allowed detecting Tau aggregates in PTX treated cells. Apparently, the massive displacement of Tau from MT, induced by taxol and the sudden increased concentration of soluble Tau triggers the aggregation. Nevertheless we cannot exclude possible PTX activities on kinases regulating Tau stability or Tau-MT interactions. Tau aggregates induced by PTX showed a fluorescence and FRET profile comparable to aggregates from synthetic Tau seeds, even if of a smaller size. Thus, PTX can be used as a simple and mechanistically well validated compound to induce Tau aggregates *in vivo*, similar to those induced by Tau seeds i) in assays to screen for inhibitors of Tau aggregation or ii) *in vivo*, to study the consequences of Tau aggregation in a well-controlled way.

Tau proteins bearing mutations involved into Tau pathology may have distinct conformations. We compared the mobility and the interaction with MTs by exploiting a panel of mutated CST sensors. The phosphorylation of S422 is considered an early marker of pathology, since it precedes the cleavage at Asp 421 and increase the aggregation propensity ^64^. Accordingly, the CST with the S422A mutation do not show any difference with respect to wt CST, as expected; on the other hand, the phospho-mimetic S422E mutation increased soluble Tau fraction. P310S or P301L Tau mutants have been shown to accelerate *in vitro* and *in vivo* the formation of aggregates ^65,66^. P301 mutated CST constructs highlight a great increase of cytoplasmic free Tau, with a doubling of the diffusive fraction for P310L (from 16% to 38%). Further analysis indicated that MT-bound P301 mutant Tau assumed a 3D-conformation not allowing an efficient FRET reaction. Thus, the distinct Tau conformation caused by the P310 mutation cold be responsible for its weaker interaction with MTs.

Finally, we tested the mutations at the AT8 epitope, that is heavily phosphorylated in NFTs and is considered a late marker of disease. AT8 phospho-mimetic CST do not differ from CST, consistently with the observation that, in Hela cells, the AT8 epitope is constitutively phosphorylated. Conversely, AT8 null mutant is tightly bound to MTs and displayed a 3D-conformation that strongly increased FRET efficiency. It is conceivable that the higher FRET could be due to both the new conformation that increase both the intramolecular and the intermolecular FRET, due to the accumulation of MT-bound Tau molecules.

There is a big interest in developing biosensors for the quantitative imaging of oligomerization and aggregation of full length Tau, in the context of living cells. Imaging of Tau aggregation in cells has been recently achieved using a biosensor system based on the aggregation-competent core Tau repeat domain fragment ^2,32^. Tau repeat domain fragments also form the basis of a sensor for the aggregation of Tau *in vitro*, at the single molecule level ^67^. In this paper we have provided an extensive validation of CST, a new conformational sensor for full length Tau in living cells. CST provides a powerful quantitative readout to acquire new information about the physiological interactions of full length Tau with MTs and its pathological aggregation responses to a variety of treatments, including chemical small molecules, synthetic Tau seeds, AD brain extracts and ApOs. Moreover, CST allowed to demonstrate the distinct effects of mutations on Tau conformations and mobility in living cells. The properties of CST make it amenable to be used both at high spatial resolution (even at single molecule resolution) in cells or in formats for a versatile set of different large-scale screenings, to advance our knowledge on Tau biology, in details such as cellular conditions or factors or molecules altering its normal interactions with MTs at the individual cell level.

## MATERIALS AND METHODS

### Chimeric constructs cloning

The cDNA encoding the Tau isoform D (383aa) has been cloned into the BspEI site of the plasmid pECFP-EYFP already available in the lab (EYFP cloned in frame with ECFP into pECFP-C1 from Clontech at the BspeI site). Both the forward and reverse cloning primers contain the RSIVT linker sequence between the BspEI site and the Tau sequence (forward primer: 5'- GTC GTT TCC GGA AGA TCT ATT GTC ACT ATG GCT GAG −3'; reverse primer: 5'- AAC GAC TCC GGA AGT GAC AAT AGA TCT CAA ACC CTG −3'). The monolabeled constructs pECFP-Tau and pTau-EYFP have been generated by subcloning the Tau cDNA into the BspEI site of plasmid pECFP-C1 and pEYFP-N1 (Clontech Laboratories, Inc., Saint-Germain-en-Laye, France)(FWD-BspEI-TAU: 5'- AAT TAT TCC GGA ATG GCT GAG CCC CGC CAG-3'; REV-BspEI-TAU: 5'- ACT TGA TCC GGA CAA ACC CTG CTT GGC CAG −3'; FWD-BspEI-RSIVT-TAU: 5'- AAT TAT TCC GGA AGA TCT ATT GTC ACT ATG GCT GAG CCC CGC CAG-3'; REV-BspEI-RSIVT-TAU: 5'- ACT TGA TCC GGA AGT GAC AAT AGA TCT CAA ACC CTG CTT GGC CAG −3'). The pTagRFP-tubulin plasmid was purchased from Evrogen (FP145).

### Cell culture, transfection and treatments

HeLa cells and immortalized hippocampal neurons HT22 were maintained in DMEM (GIBCO) supplemented with 10% FCS; SHSY-5Y cells were maintained in DMEM-F12 (GIBCO) supplemented with 10% FCS and differentiated adding 10uM RA for 5 days. The day before the experiment cells were seeded at 10x10^4^ cells per well in six-well plates or in wilco dishes (Willcowells). The lipofection was carried out with Effectene (QIAGEN) according to manufacturer’s instructions. Cells have been treated with TNFα (Peprotech, 20 ng/ml) 24h post-transfection. Cells have been treated with 1uM Nocodazole (Sigma) for 20 min, or with 1uM Colchicine (Sigma) for 24 hours. Cells have been treated with 10uM Staurosporin (Sigma) or with 1uM Paclitaxel (Sigma), or with 1uM Epothilone D (Santa Cruz Biotechnology). AD brain lysates have been provided by Abcam (ab29969, ab29971) and cells have been treated with 10ug. Tau seeds have been produced by *in vitro* oligomerization. The reaction has been performed with 4mg/ml Tau (297-391)4R in PBS pH 7,2 and 100uM Heparin, incubated for 4 days and centrifuged at 100000xg for 1,5h. ABOs have been collected from 7PA2 cells conditioned medium as described by ^46^, and 500ul of AβOs have been added to 1 ml of complete medium for 96h.

### Western blot and Immunostaining

Cells extracts were prepared in lysis buffer supplemented with protease and phosphatase inhibitors by lysis on ice for 30 minutes. Total proteins were separated by 10% or 8% SDS-PAGE and electro-blotted onto nitrocellulose membranes Hybond-C-Extra (Amersham Biosciences). Membranes were blocked with 5% skimmed milk powder in TBS containing 0.1% Tween 20.

Primary antibodies for Western blot analysis were: anti Tau5, anti Tau-C3, anti Tau12, anti GFP ( Abcam), anti GAPDH (Fitzgerald), anti DC25 from M. Novak. Secondary antibodies for western blot analysis were HRP-conjugated anti-mouse or anti-rabbit or anti-goat IgG were purchased from Santa Cruz Biotechnology, Inc. For immunofluorescence experiments cells were fixed with cold ethanol for 5 minutes at 4°C. After permeabilization with PBS 1X containing 0.2% Triton-X100 for 10 minutes, samples were blocked with 2% bovine serum albumin. The slides were incubated with the primary antibody 1 hour at room temperature and with secondary antibodies fluorophore-conjugated 1 hour 1at room temperature. Slides were mounted with Vectashield mounting medium (Vector Laboratories). Primary antibodies used for immunofluorescence were: anti Tau-13 (SCBT) and anti-tubulin (Abcam). Secondary antibodies used for immunofluorescence were: anti-rabbit or anti-mouse conjugated with Alexa-594, Alexa-633 (Molecular Probes, Eugene, OR)

### Preparation of Sarkosyl-insoluble Tau

Sarkosyl-insoluble Tau was prepared by diluting cells lysate in 0.5 ml of A68 buffer (10 mm Tris-HCl, pH 7.4, 0.8 m NaCl, 1 mm EGTA, 5 mm EDTA, 10% sucrose), supplemented with protease and phosphatase inhibitors, followed by centrifugation at 13,000 × g at 4 °C for 20 min. Samples were then incubated with 1% sodium lauroyl sarcosinate (Sarkosyl, Sigma) at room temperature for 1 h in a flat rotating shaker at 700 rpm followed by centrifugation at 100,000 × g at 4 °C for 1 h. The pellets were washed and resuspended in 50 mm Tris-HCl, pH 7.4. Samples were used immediately or snap-frozen in liquid nitrogen and stored at −80 °C.

### Image acquisition and analysis

Images were acquired with the TCS SL laser-scanning confocal microscope (Leica Microsystems, Milan, Italy) equipped with galvanometric stage using a 63X/1.4 NA HCX PL APO oil immersion objective. An heated and humidified chamber mounted on the stage of the microscope was used for live imaging experiments in order to maintain a controlled temperature (37°C) and CO2 (5%) during image acquisition. An Ar laser was used for ECFP (l = 458 nm), EYFP (l = 514 nm, RFP (l=453 nm) and Alexa 633 (l=633 nm). For the quantification of morphological parameters such as the total filament length and the number of branching points the filament tracer option of the Imaris Bitplane software has been exploited. In detail, these two parameters are deduced by a software plugin that, based on connectivity and fluorescence intensity, automatically detects and segments filamentous structures revealing information about the topology of filaments as the sum of the lengths of all lines and the number of branching points within the filament.

### FRET and FRAP experiments

For sensitized emission FRET experiments, each image was recorded in a spectral mode, by selecting specific regions of the emission spectrum. In particular, the donor ECFP was excited at 458 nm and its fluorescence emission was collected between 470 nm and 500 nm (donor channel) and between 530 nm and 600 nm (FRET channel). The acceptor EYFP was excited at 514 nm and its fluorescence emission was collected between 530 nm and 600 nm (acceptor channel). The donor and acceptor fluorophores were excited sequentially. The ImageJ software was used for images analysis and FRET quantification. Briefly, FRET images were corrected from cross-talk between donor and acceptor channel using Youvan’s method^68^: F_index_=I_FRET_-A X I_D_-B X I_A_, where I_FRET_, I_D_ and I_A_ are the images of the sample in the FRET, donor and acceptor channel, respectively, after background subtraction and A and B are the fraction of the donor and acceptor leak-through into the FRET channel, respectively. A and B parameters have been evaluated in cells expressing only the donor and only the acceptor (A=0,1 and B=0,25). In our experimental condition, no leak-through signal from ECFP and EYFP channel or viceversa was observed. Normalized FRET was performed using: NFRET=F_index_/√(I_D_ X I_A_) (REF). NFRET intensities images were represented in false-color scale. FRAP experiments were performed by using the FRAP module coupled to the confocal microscope and consists of three different phases: 1) a pre-bleach phase, in which 10 frames of 512*512 pixel images at a 1000Hz have been recorded in order to define the initial level of fluorescence intensity; 2) a photobleaching phase, in which a selected circular ROI with a radius of 2um in the cytoplasm of the cell was excited at higher laser power (50% for EYFP) for 5 frames at 1000Hz; 3) a post-bleaching phase, in which 120 images have been recorded in order to follow the recovery of the fluorescence intensity in the selected ROI. Fluorescence recovery was extracted from images of the bleached ROI and subjected to the following manipulation steps:1) background subtraction; 2)first normalization to the initial pre-bleach value of fluorescence intensity; 3) correction for the fluorescence loss; 4) additional normalization to set the first post-bleach point to zero. At least 30 separate FRAP experiments for each sample has been performed. FRAP recovery curves have been fitted by a two phase exponential association function (OriginLab).

**Figure 4.**
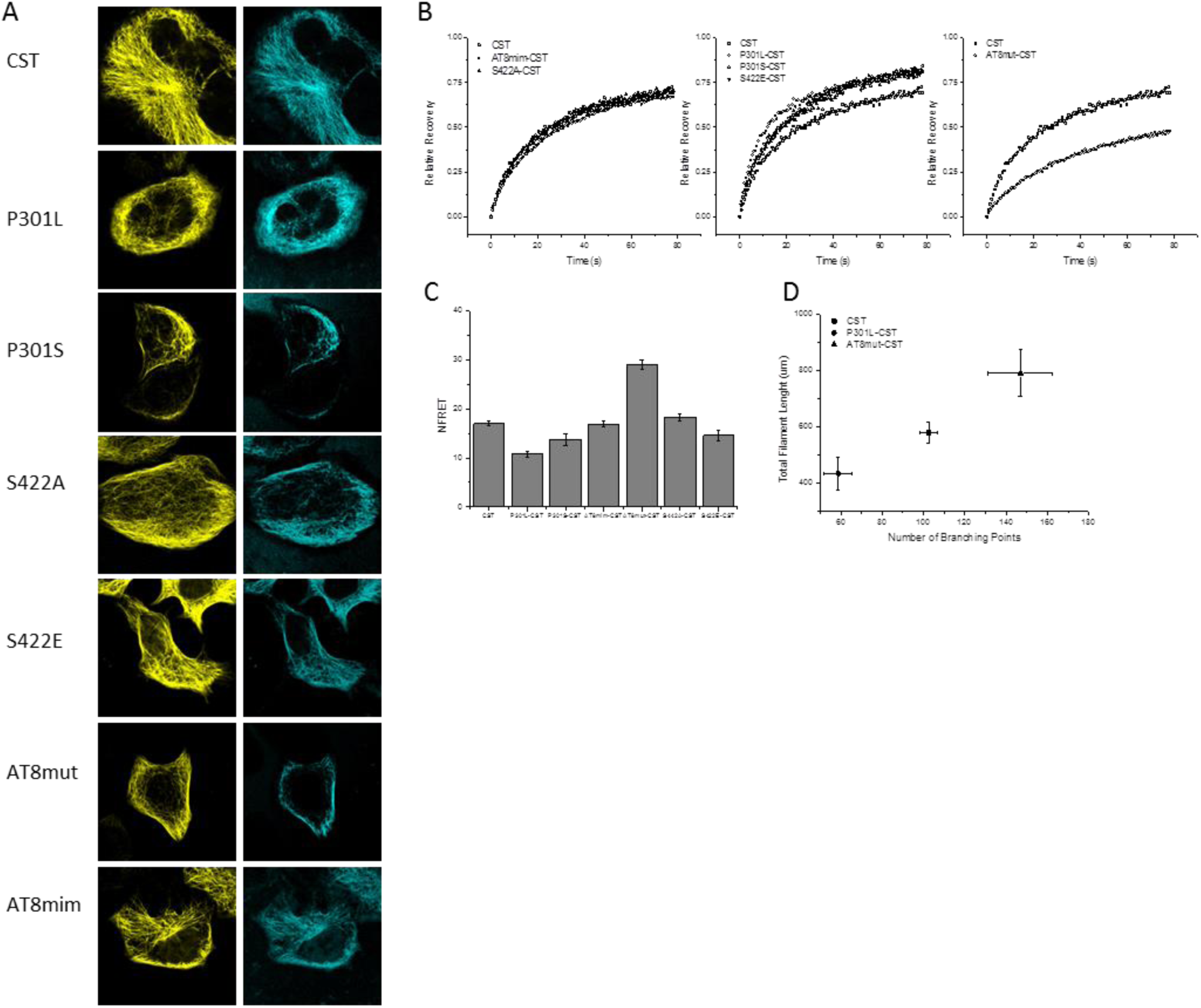
FRAP and FRET analysis of mutant CST. A. Imaging of acceptor (yellow) and donor (blue) channel from CST mutated constructs in living cells. B. FRAP recovery curves of mutated CST constructs. C. Normalized FRET values measured on fluorescent MTs in reporter cells. D. Morphological analysis of Tau network complexity in reporter cells expressing CST or P301L-CST or AT8mut-CST.

## FIGURE LEGENDS

**Suppl.fig.1**. A. Schematic representation of antibodies antigens on CST construct. B. Western blot of total extracts from HeLa transfected cells. C. Anti Tau13 antibody immunofluorescence in HeLa cells (green). D. CST imaging in SHSY5Y cells (yellow). E. Endogenous Tau immunofluorescence with Tau13 antibody in differentiated SHSY5Y cells (red).

**Suppl. fig.2.** Fluorescence intensity distribution of reporter cells expressing monolabeled Tau constructs with a comparable concentration of Tau and half concentration of fluorophores with respect to CST (CFP-Tau+Tau-YFP)/2, monolabeled Tau constructs with a double amount of Tau and same amount of fluorophores with respect to CST (CFP-Tau+Tau-YFP).

**Suppl.fig.3.** Imaging of the acceptor channel (yellow), FRET (purple) and tubulin immunofluorescence (red) of reporter cells treated with colchicine or nocodazole. Western blot analysis of total extracts from untreated and treated cells.

**Suppl. fig. 4** Colocalization between CST aggregates (green) and K114 staining (red) in PTX treated cells. White arrows highlight colocalization spots.

**Suppl fig. 5** MT network is not altered after treatment with: Tau seeds, human AD brain lysates, AD11 brain lysates, PTX, EpoD, AβOs. Imaging of CST (yellow) and immunofluorescence of tubulin (red).

**Suppl.fig.6.** CST imaging in HT22 cells treated with Tau seeds, human AD brain lysates and AβOs. Reporter cells have been treated for 48h with Tau seeds or for 96h with cleared human brain lysates from healthy donors or from AD donors or with supernatant containing AβOs. Donor channel (blue) and acceptor channel (yellow) have been acquired.

**Suppl. Table 1** FRAP values of wt and mutated CST

